# Improving deep learning-based protein distance prediction in CASP14

**DOI:** 10.1101/2021.02.02.429462

**Authors:** Zhiye Guo, Tianqi Wu, Jian Liu, Jie Hou, Jianlin Cheng

## Abstract

Accurate prediction of residue-residue distances is important for protein structure prediction. We developed several protein distance predictors based on a deep learning distance prediction method and blindly tested them in the 14th Critical Assessment of Protein Structure Prediction (CASP14). The prediction method uses deep residual neural networks with the channel-wise attention mechanism to classify the distance between every two residues into multiple distance intervals. The input features for the deep learning method include co-evolutionary features as well as other sequence-based features derived from multiple sequence alignments (MSAs). Three alignment methods are used with multiple protein sequence/profile databases to generate MSAs for input feature generation. Based on different configurations and training strategies of the deep learning method, five MULTICOM distance predictors were created to participate in the CASP14 experiment. Benchmarked on 37 hard CASP14 domains, the best performing MULTICOM predictor is ranked 5th out of 30 automated CASP14 distance prediction servers in terms of precision of top L/5 long-range contact predictions (i.e. classifying distances between two residues into two categories: in contact (< 8 Angstrom) and not in contact otherwise) and performs better than the best CASP13 distance prediction method. The best performing MULTICOM predictor is also ranked 6th among automated server predictors in classifying inter-residue distances into 10 distance intervals defined by CASP14 according to the F1 measure. The results show that the quality and depth of MSAs depend on alignment methods and sequence databases and have a significant impact on the accuracy of distance prediction. Using larger training datasets and multiple complementary features improves prediction accuracy. However, the number of effective sequences in MSAs is only a weak indicator of the quality of MSAs and the accuracy of predicted distance maps. In contrast, there is a strong correlation between the accuracy of contact/distance predictions and the average probability of the predicted contacts, which can therefore be more effectively used to estimate the confidence of distance predictions and select predicted distance maps.

## 1 Introduction

Accurate prediction of inter-residue distances (or its simplified representation – inter-residue contacts) is critical for template-free (ab initio) tertiary structure prediction, i.e., predicting the structure of a protein without using any known structure as templates (Kryshtafovych, et al., 2019). The predicted inter-residue distances can be translated into tertiary structures by off-shelf tools such as trRosetta (Yang, et al., 2020), CONFOLD2 (Adhikari and Cheng, 2018) built on top of CNS (Brünger, et al., 1998), and DMPfold (Greener, et al., 2019). In the 2018 CASP13 experiment, the top-ranked methods (Hou, et al., 2019; Kandathil, et al., 2019; Senior, et al., 2020; Xu and Wang, 2019; Zheng, et al., 2019) all used distance or contact predictions to guide template-free (FM) structure modeling to achieve significant success. Since then, the inter-residue distance prediction has become a focal point of protein structure prediction.

In the last several years, the advances in protein distance/contact prediction were mostly driven by two technologies: the residue-residue co-evolutionary analysis (Ekeberg, et al., 2013; Kamisetty, et al., 2013; Seemayer, et al., 2014) for generating informative features for prediction and various deep learning methods (Goodfellow, et al., 2013; He, et al., 2016) for effectively extracting protein distance/contact patterns from the features. Since classifying the distances between residues into multiple distance intervals (commonly called distance prediction) can provide more detailed information about residue-residue distances than classifying them into two binary categories – in contact or not in contact (commonly called contact prediction), recent methods such as AlphaFold and RaptorX focus on the distance prediction. The multi-classification or binary classification of distances produces a multi-class or binary-class distance probability map. Most recently, some methods such as DeepDist (Wu, et al., 2020) were developed to predict real-value inter-residue distances using deep learning regression methods, in addition to classifying the distances into multiple distance intervals. Moreover, the attention mechanism that can pick up relevant signals anywhere in the input features was also applied to predict protein contacts and explain the predictions (Chen, et al., 2020). In the CASP14 experiment, the attention mechanism was also used by AlphaFold2, tFold, and our MULTICOM distance predictors to improve distance prediction.

In this work, we describe the design and implementation of our MULTICOM distance predictors based on our DeepDist2 distance prediction method and analyze their results and performance in CASP14. Following the CASP14 norm, the analysis is focused on hard template-free modeling (FM) target domains instead of template-based modeling (TBM) domains that have recognizable known template structures in the Protein Data Bank (PDB) (Berman, et al., 2000). The FM/TBM domains that might have very weak templates that cannot be recognized by existing sequence alignment methods are also used in the evaluation.

## 2 Materials and Methods

The overall pipeline of the MULTICOM distance predictors based on our latest deep learning method – DeepDist2 is shown in **Fig.1**. Three methods are used to generate multiple sequence alignments (MSAs) for a target protein in parallel, including our in-house tool – DeepAln (Wu, et al., 2020), DeepMSA (Zhang, et al., 2019), and HHblits (Remmert, et al., 2012). DeepAln and DeepMSA are also used in the original DeepDist method. In CASP14, MULTICOM predictors added the HHblits search against the Big Fantastic Database (BFD) (Steinegger, et al., 2019) (denoted as HHblits_BFD) to generate MSAs when the number sequences in MSAs generated by DeepAln and DeepMSA was less than 10L (L: sequence length).

**Fig. 1.**
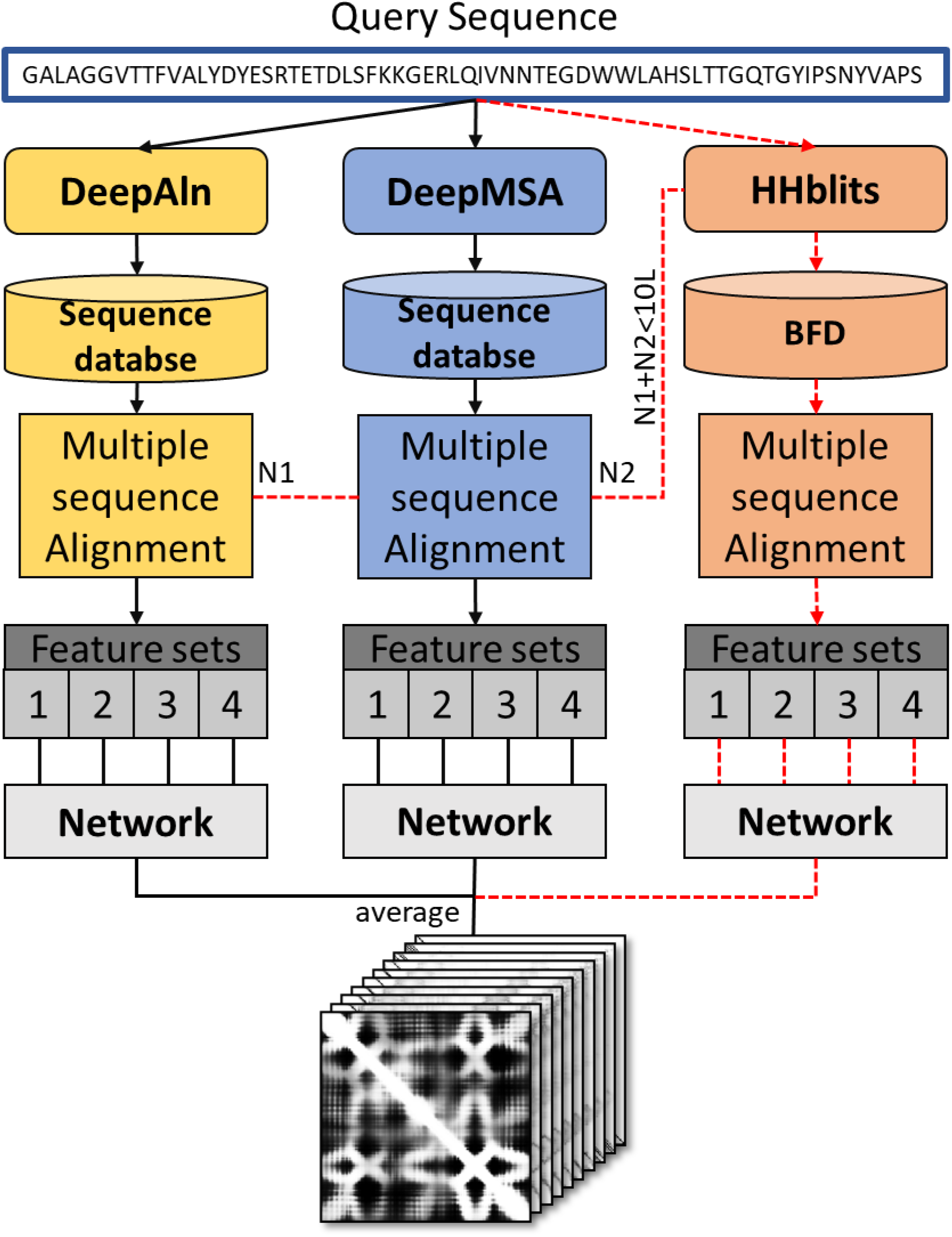
The overall pipeline of the MULTICOM distance predictors based on DeepDist2. The two data flows (branches) applied to all the targets are connected by the black solid line, while the optional flow (branch) is connected by the red dotted line, which is only invoked when the MSAs are produced by DeepMSA and DeepAln are not sufficiently deep. Each flow (branch) produces four sets of features (COV_Set, PRE_Set, PLM_Set, and OTHER_Set; see details in Section 2.2), each of which is used as input for a deep network to predict a distance map. The four distance maps predicted from the four sets of features of each branch are averaged as the predicted distance map of the branch. The final prediction is the average of the predicted distance maps of the first two or all the three branches.

Each MSA is used to produce multiple co-evolutionary features such as covariance matrix (Jones and Kandathil, 2018), precision matrix (Li, et al., 2019), and pseudolikelihood maximization matrix (Seemayer, et al., 2014). The quality of the co-evolutionary features depends on the depth of MSA (i.e. the number of sequences) as well as the quality of the MSA (e.g., the proportion of true homologous sequences in MSA). For instance, when the number of effective sequences (Neff) in an MSA is too small, the co-evolutionary scores tend to be noisy and less informative (Wu, et al., 2020). To complement the co-evolutionary features, the non-coevolutionary features such as position-specific scoring matrix (PSSM) generated by PSI-BLAST (Bhagwat and Aravind, 2007) and secondary structures are also used.

Different kinds of co-evolutionary features are combined with non-co-evolutionary features to generate the four sets of features (COV_Set, PRE_Set, PLM_Set, and OTHER_Set; see details in Section 2.2). Each of four sets of features derived from the same MSA is used by a deep residual network with a channel-wise attention mechanism to predict a distance map. The average of the four predicted distance maps is the predicted distance map for the MSA. Different from DeepDist that uses four different deep architectures for different sets of features, DeepDist2 uses the same network architecture for all the feature sets. For most CASP14 targets, the distance maps predicted from the features generated from DeepAln’s MSA and DeepMSA’s MSA were averaged as the final prediction. When the number of sequences in the combination of MSAs generated by DeepAln and DeepMSA was less than 10 L, the distance map predicted from the MSA of HHblits_BFD was averaged with the distance maps predicted from MSAs of DeepAln and DeepMSA as the final prediction.

Based on the same protocol above, four automated MULTICOM distance predictors (MULTICOM-CONSTRUCT, MULTICOM-AI, MULTICOM-HYBRID, MULTICOM-DIST were trained with different labelings of distance intervals. MULTICOM-DEEP used the average of the four predictors as its prediction. The distance intervals (or bins) of MULTICOM-CONSTRUCT are 0 to 4 Å, 4 to 6 Å, 6 to 8 Å, …, 18 to 20 Å, and > 20 Å. MULTICOM-DIST uses 42 bins, i.e. dividing 2 to 22 Å into 40 bins with a bin size of 0.5 Å, plus 0 – 2 Å bin and > 22 Å bin. MULTICOM-HYBRID shares the same distance segmentation strategy as MULTICOM-DIST, except that it starts with an interval 0 – 3.5 Å and its last interval is set to > 19 Å. MULTICOM-AI has 37 equally spaced intervals of 0.5 Å between 0 to 20 Å and the > 20 Å interval. Though the predicted multiclass distance prediction maps of the five predictors are based on the different distance intervals, they are converted into the 10-bin classification maps required by CASP14. The 10 bins defined CASP14 are bin1: d≤4Å, bin2: 4<d≤6Å, bin3: 6<d≤8Å, …, bin10: >20Å, which are the same as MULTICOM-CONSTRUCT.

### 2.1 Deep residual neural networks with channel-wise attention mechanism for inter-residue distance prediction

The architecture of the deep residual network with the attention mechanism is shown in **Fig.2**. The input features (a tensor of L * L * N dimension; L: sequence length; N: number of channels) are first fed into an instance normalization layer (Ulyanov, et al., 2016), followed by a convolutional layer and a Maxout layer (Goodfellow, et al., 2013). The convolutional layer reduces the number of channels to 128 and then the Maxout layer halves it to 64.

**Fig. 2.**
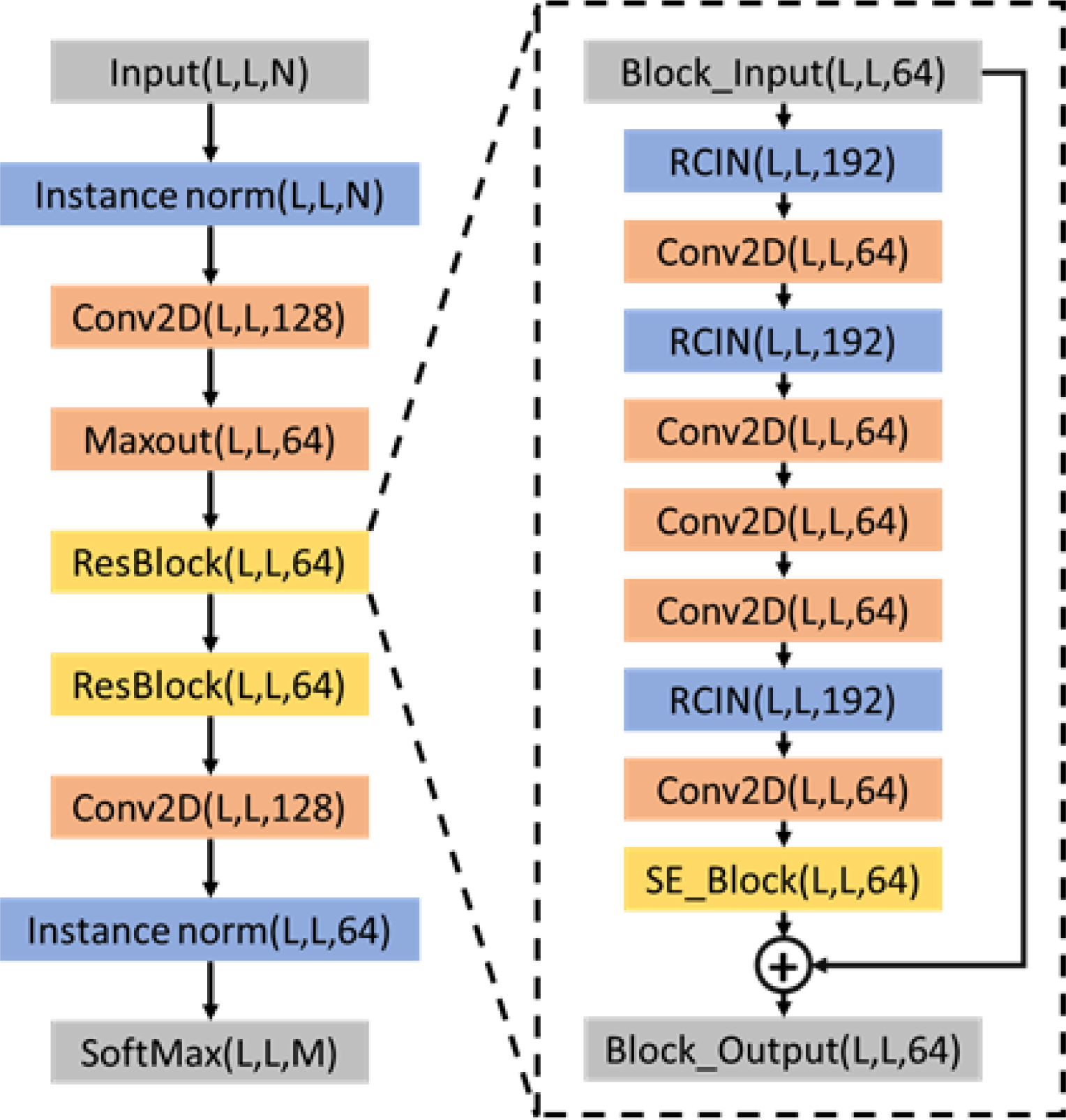
The architecture of the deep residual network of DeepDist2 used by MULTICOM distance predictors. L is the length of the input sequence. N is the number of channels of the input features (i.e. 483 for COV_set, 482 for PLM_set, 484 for PRE_set, and 47 for OTHER_set).

Following the Maxout layer are 20 residual blocks with the same input and output dimension of 64. Each residual block starts with a normalization block (called RCIN) that includes three different kinds of normalization layers and one ReLU (Nair and Hinton, 2010) activation function. The three normalization layers of RCIN are row normalization layer (RN), column normalization layer (CN) (Mao, et al., 2019), and instance normalization (IN) layer. The output of the three normalization layers is concatenated as input for a ReLU activation function. Through this operation, the information in multiple directions can be effectively integrated to better capture contacts/distances between residues. The RCIN block is followed by a convolutional layer, an RCIN block, three convolutional layers, an RCIN block, and a convolutional layer. The final part of the residual block is the squeeze-and-excitation block (SE) (Hu, et al., 2018), which is a channel-wise attention method popular in the computer vision field. This block has good adaptability and can be embedded into different deep network architectures. It has two parts: one is the squeeze operation that can collect the global information between all the feature channels and another is the excitation operation that can boost the impact of relevant features by two fully connected layers with the ReLU activation function. The SE block recalibrates the feature channels through learning so that the network can assign more attention to more essential feature channels. We apply a softmax activation to classify inter-residue distances between residues into multiple intervals (bins), i.e. predict the probability distribution of inter-residue distances.

### 2.2 Multiple sequence alignments and input features

DeepAln and DeepMSA use HHblits and jackhmmer to search several protein sequence datasets to generate MSAs (Wu, et al., 2020). During the CASP14 experiment, all the databases (i.e. UniRef90 (2020-04) (Mirdita, et al., 2017), Uniclust30 (2020-03), Metaclust50 (2018-06) (Steinegger and Söding, 2018), Myg_UniRef100) used for MSA generation were updated to their latest version. The BFD used by HHblits search was released by March 2019.

The residue-residue co-evolutionary features including covariance matrix (COV), precision matrix (PRE), and pseudolikelihood maximization matrix (PLM) calculated from MSAs are two-dimensional (2D) features with multiple channels and have the dimension of L×L×441. PSSM generated from PSI-BLAST search against UniRef90 is also a useful feature. Other features like the Pearson’s correlation between columns of PSSM, the co-evolutionary contact scores produced by CCMpred, the Shannon entropy sum, mean contact potential, normalized mutual information, and mutual information from DNCON2 are generated. These features are combined to generate four sets of features as follows. COV_Set includes COV, PSSM, Pearson correlation, and CCMpred contact scores; PLM_Set contains PLM, PSSM, Pearson’s correlation; PRE_Set has PRE, PSSM, and entropy scores (joint entropy, Shannon entropy sum); and OTHER_Set has PSSM, CCMpred contact scores, Pearson correlation, solvent accessibility, mean contact potential, normalized mutual information, and mutual information.

### 2.3 Datasets and evaluation metrics

11,234 proteins were used to train the MULTICOM distance predictors. The proteins in the training dataset have less than 25% sequence identity with the proteins in the three test datasets: 43 CASP13 FM and FM/TBM domains, 37 CASP12 FM domains, and 268 CAMEO targets (released between 08/31/2018 and 08/24/2019). The predictors were trained and internally tested on the test datasets before they were blindly tested in CASP14 from May to July 2020.

Our evaluation of the MULTICOM distance predictors was based on 37 hard FM and FM/TBM domains of CASP14 (i.e. 23 FM domains and 14 FM/TBM domains). To be consistent with the analysis of CASP14, the evaluation is carried out at the domain-level. The distance predictions are evaluated by three metrics: (1) the precision of top L/5, L/2, or L long-range contact prediction after the multi-class distance predictions are converted to binary contact predictions at 8 Å threshold (L: sequence length), (2) mean absolute error (MAE) between predicted distances and true distances; and (3) the average precision, recall, and F-measure of multi-classification of distances between long-range residue pairs over 10 distance bins.

Two residues are considered in contact if the distance between their β-carbon atoms (α-carbon for the glycine amino acid) is less than 8 Å. A contact map can be obtained by summing up the probability values of the intervals within 0-8 Å in a predicted multi-classification distance map. We use ConEVA (Adhikari, et al., 2016) to calculate the precision of predicted contacts. The CASP14’s assessment results at https://predictioncenter.org/casp14/rrc_avrg_results.cgi are also used. A contact is considered long-range contact if the sequence separation between the two residues is >= 24 residues, medium-range if the sequence separation is within [12, 23], and short-range if the sequence separation is within [6, 11]. In this study, the evaluation is mostly focused on long-range residue-residue contact/distance predictions according to the CASP norm.

The real-value distance between two residues is estimated as the sum of the mean distance of each interval times the predicted probability of the interval (i.e. the weighted average). Because large distances contribute little to tertiary structure prediction, only predicted distances less than 16 Å are used for the MAE evaluation. The standard deviation of the MAE is also calculated. When the MAE is close, the smaller standard deviation is preferred.

For the multi-classification prediction, we apply the precision (denoted as Precision_m), recall (denoted as Recall_m) to evaluate the multi-classification of distances between long-range residue pairs. The precision and recall of each distance bin for a target are calculated first. The precision and recall of multiple distance bins is the arithmetic average of precision and recall of each bin over all the bins. Therefore, the final precision and recall (Precision_m and Recall_m) can evaluate the accuracy of the overall performance of multi-classification of distances for a target. We only calculate the precision and recall of the multi-classification prediction of the distances between long-range residue-residue pairs. The F1-measure is the geometric mean of Precision_m and Recall_m.

## 3 Results

### 3.1 Overall performance of distance prediction in CASP14

In this study, we only compare CASP14 server predictors, excluding CASP14 human predictors that had more prediction time and might use some server predictions as input. The performance of the top 20 out of 30 CASP14 automated server predictors on 22 FM domains in terms of precision of top L/5 long-range contact predictions (called top L/5 precision) is shown in **Table 1**. The top L/2 precision of the predictors is also reported in the table. The result was directly compiled from the evaluation data at the CSAP website after excluding human distance predictors. Our best server predictor MULITCOM-CONSTRUCT has a top L/5 precision of 64.99% and is ranked no. 5 after TripletRes from Zhang Group and three tFold servers (tFold-CaT, tFold-IDT, and tFold) from tFold Group. Other MULTICOM predictors are also ranked among the top 20. Moreover, the top L/5 (or L/2) precision of the MULTICOM predictors is higher than RaptorX – the best contact predictor in CASP13, showing that multiple predictors including ours in the CASP14 experiment improve over the best CASP13 contact predictor.

**Table1.**
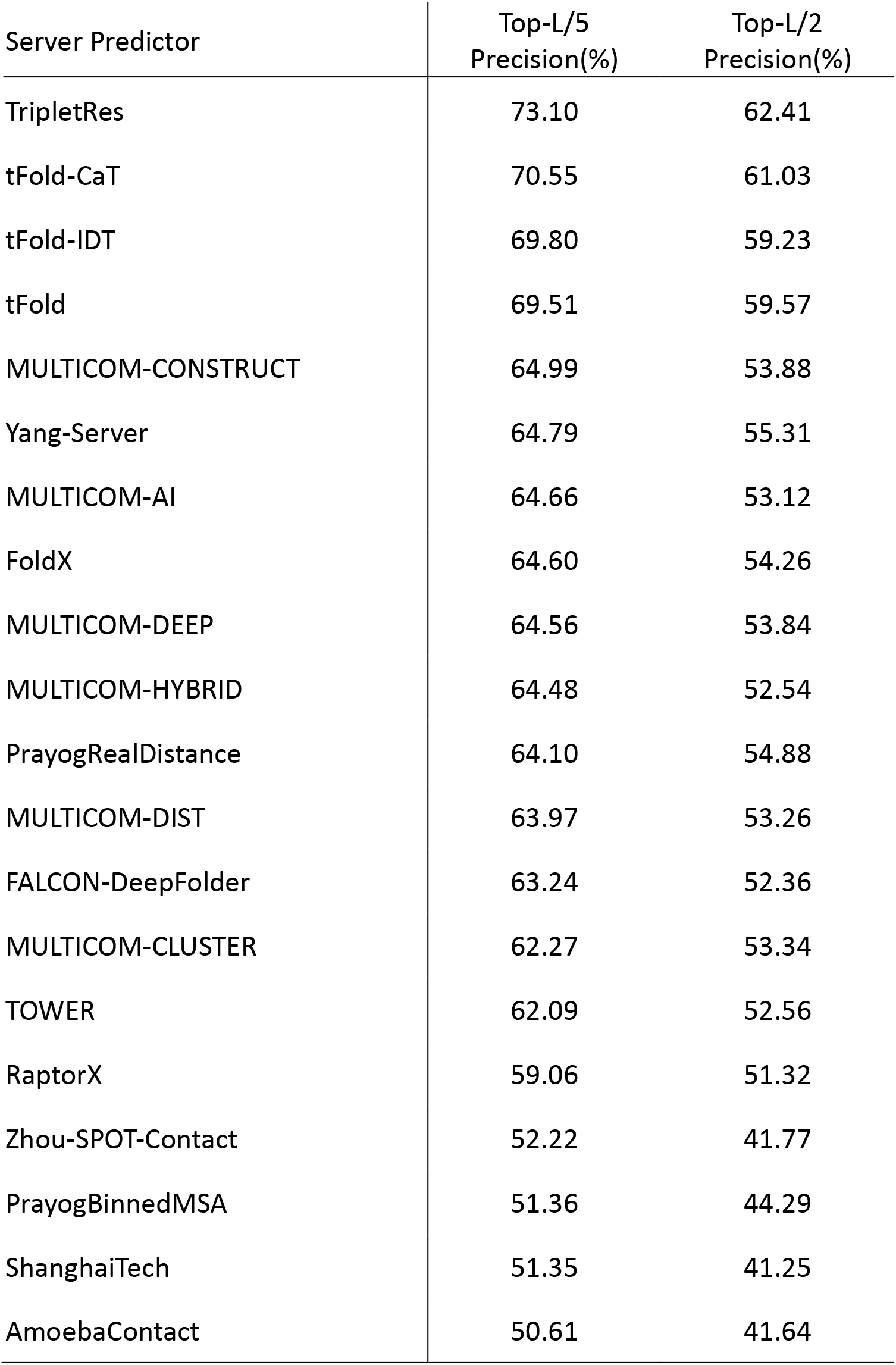
The performance of top 20 server predictors on CASP14 37 FM and FM/TBM domains in terms of the precision of top L/5 and top L/2 long-range contact precisions. The predictors are ranked by the top L/5 precision.

Among the 30 server predictors, 19 of them submitted multi-class distance predictions, while the rest only submitted binary contact predictions. **Table 2** reports the precision (Precision_m), recall (Recall_m), and F1-measure of multi-classification distance prediction of the 19 predictors. Our best server predictor MULTICOM-DIST is no.6 after TripletRes from Zhang group, three tFold servers (tFold-CaT, tFold-IDT, and tFold) from tFold group, and Yang-Server from Yang group in terms of F1-measure.

**Table2.** The performance of CASP14 server predictors on 22 FM domains in terms of precision_m, recall_m, and F1 measure of multi-class distance prediction. The server predictors are ranked by the F1 measure.

| Server Predictor | Precision_m(%) | Recall_m(%) | F1 measure |
| --- | --- | --- | --- |
| TripletRes | 31.90 | 24.24 | 0.2493 |
| tFold-IDT | 29.03 | 20.71 | 0.2179 |
| tFold-CaT | 29.92 | 20.03 | 0.2115 |
| tFold | 33.25 | 18.61 | 0.1955 |
| Yang-Server | 26.55 | 18.22 | 0.1857 |
| MULTICOM-DIST | 28.22 | 17.70 | 0.1847 |
| FoldX | 28.88 | 18.07 | 0.1832 |
| MULTICOM-CONSTRUCT | 27.12 | 17.64 | 0.1828 |
| MULTICOM-DEEP | 27.98 | 17.52 | 0.1815 |
| TOWER | 28.35 | 17.81 | 0.1792 |
| MULTICOM-AI | 27.72 | 17.37 | 0.1791 |
| MULTICOM-HYBRID | 25.85 | 17.10 | 0.1764 |
| Yang_FM | 27.35 | 16.94 | 0.1762 |
| FALCON-DeepFolder | 28.00 | 16.81 | 0.1733 |
| Kiharalab_Z_Server | 21.56 | 15.63 | 0.1576 |
| RaptorX | 26.37 | 13.61 | 0.1428 |
| Zhou-SPOT-Contact | 23.27 | 13.14 | 0.1374 |
| RBO-PSP-CP | 15.97 | 13.03 | 0.1180 |
| ProdGAN_Gonglab | 6.61 | 9.50 | 0.0422 |

The detailed results of the MULTICOM distance predictors (precision of top L/5, L/2, L long-range contact predictions, the mean absolute error and standard deviation of long-range distance predictions, and the precision_m and recall_m of multi-classification of distances) on 37 FM and FM/TBM domains are reported in **Table 3**. The MULTICOM distance predictors have similar performance. MULTICOM-CONSTRUCT performs best in terms of contact precision, MULTlCoM-AI has the lowest MAE, and MULTICOM-DEEP has the highest multi-classification precision.

**Table 3.** The performance of distance predictions of MULTICOM server predictors on the 37 CASP14 FM and FM/TBM domains. Bold denotes the highest value.

| | Top-L/5<br>(%) | Top-L/2<br>(%) | Top-L (%) | MAE±std | Precision_<br>m (%) | Recall_m<br>(%) |
| --- | --- | --- | --- | --- | --- | --- |
| MULTICOM-AI | 64.66 | 53.12 | 40.75 | **2.631**±2.10 | 33.14 | 21.94 |
| MULTICOM-CONSTRUCT | **64.99** | **53.88** | 41.25 | 2.665±2.11 | 32.08 | **22.02** |
| MULTICOM-DEEP | 64.56 | 53.84 | **41.32** | 2.658±2.11 | **33.25** | 21.88 |
| MULTICOM-DIST | 63.97 | 53.26 | 40.35 | 2.684±2.11 | 32.69 | **22.02** |
| MULTICOM-HYBRID | 64.48 | 52.54 | 40.55 | 2.743±2.15 | 30.69 | 21.32 |

### 3.2 Comparison of different MSAs for distance prediction

The performance of deep learning distance predictors depends on the quality of the input features, particularly the most important co-evolutionary features whose quality is largely determined by the depth and quality of MSAs (Wu, et al., 2020).

The depth of an MSA is usually measured by the number of effective sequences (Neff) in the MSA. Here we use the performance of MULTICOM-CONSTRUCT with three kinds of MSAs on the 37 FM and FM/TBM domains to compare their performance in distance prediction. **Table 4** shows the performance of the long-range distance prediction of MULTICOM-CONSTRUCT with MSAs of DeepAln, DeepMSA, and HHblits_BFD according to multiple metrics, including Top L/2 and Top L precisions of long-range contact predictions, mean absolute error of long-range predicted distances < 16 Å (MAE_16) and their standard deviation (STD_16), the accuracy and recall of multi-classification of distances (Precision_m and Recall_m). HHblits_BFD performs best among the three according to all the metrics, DeepMSA works better than DeepAln. For instance, the top L/2 precision of HHbits_BFD is 51.33%, higher than DeepMSA’s 46.18% and DeepAln’s 43.87%. The reason is that the BFD database (released in April 2019) contains the hidden Markov model (HMM) profiles for both proteins in UniProt and the metagenomics databases, which enables HHblits to generate high-quality alignments with the sequences in the databases. In contrast, DeepMSA or DeepAln uses HHblits to search the HMM profiles in UniProt and Jackhmmer to the sequences in the metagenomics database. Because Jackhammer’s alignment quality and sensitivity are lower than HHblits, even though DeepMSA and DeepAln search a target against a newer version of UniProt and metagenomics databases than the BFD database, the quality gain of HHblits search on the BFD still outweighs the increase of the size of databases used by DeepMSA and DeepAln, leading to the better distance predictions with HHblits_BFD.

**Table 4.** Distance prediction performance of MULTICOM-CONSTRUCT with MSAs of DeepAln, DeepMSA, and HHblits_BFD (HHblits search on the BFD database) on the 31 FM and FM/TBM CASP14 targets. Five different metrics are used to evaluate the long-range distance predictions.

|  | Top-L/2<br>Precision (%) | Top-L<br>Precision (%) | MAE_16(Å) | Precision_m<br>(%) | Recall_m (%) |
| --- | --- | --- | --- | --- | --- |
| DeepAln | 43.87 | 33.08 | 4.6733±7.87 | 29.34 | 18.51 |
| DeepMSA | 46.18 | 35.38 | 4.4538±7.80 | 30.49 | 19.25 |
| HHblits_BFD | 51.33 | 38.37 | 4.211±7.75 | 35.42 | 20.53 |

To further quantitatively analyze the impact of different MSA generation pipelines on the performance of the distance prediction, we study the relationship between the accuracy of distance prediction and the logarithm of the number of effective sequences (Neff) in the MSAs generated by DeepAln, DeepMSA, and HHblits_BFD in **Figure 3**. Because our automatic domain parsing did not predict domains accurately in many cases during CASP14 and therefore their predicted domain boundaries are different from the ground truth, here we only analyze the 31 full-length targets in which the 37 FM and FM/TBM domain are located. The Neff and prediction accuracy are calculated on the 31 full-length targets.

**Fig. 3.**
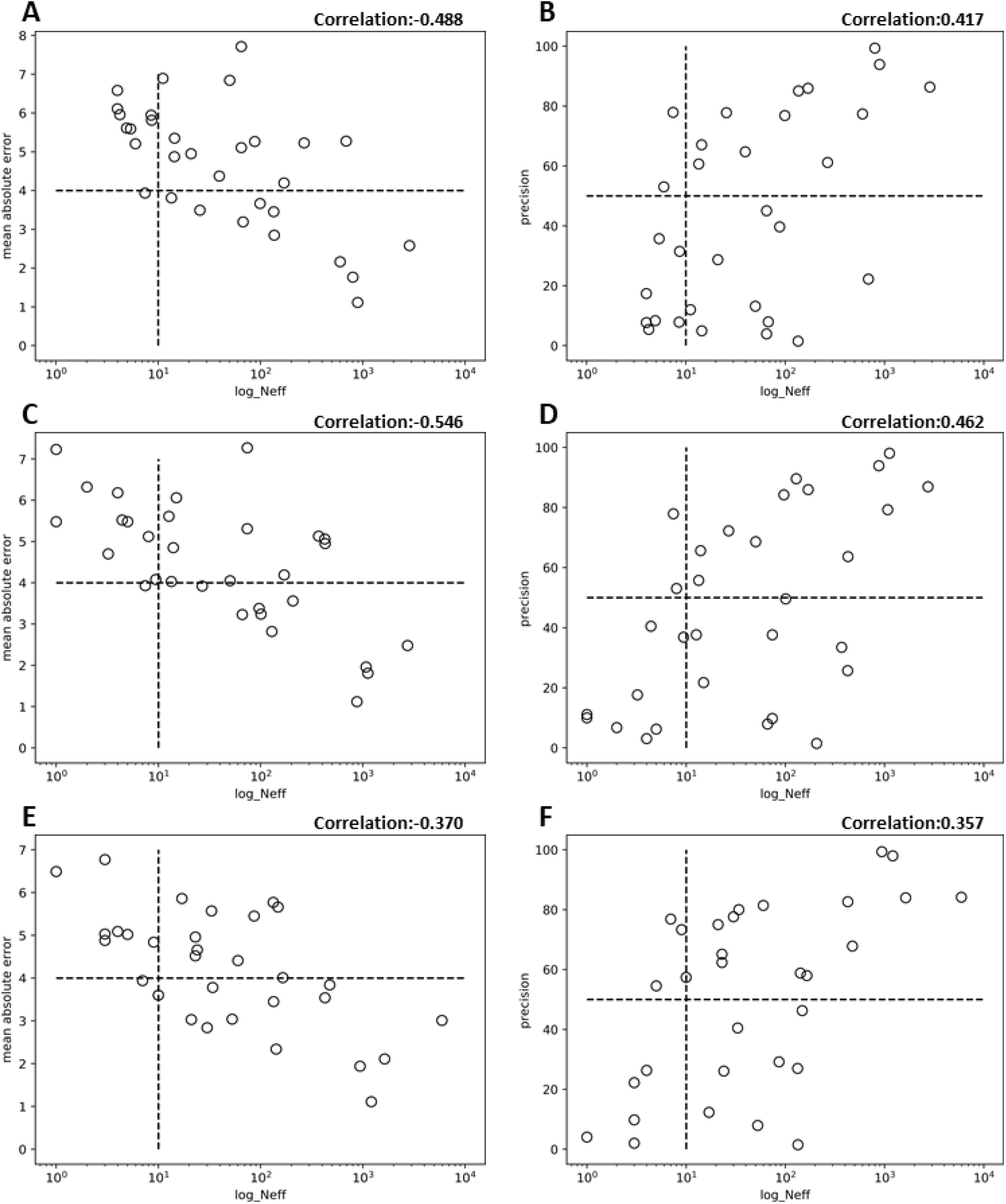
Illustration of the effect of different multiple sequence alignment generation pipelines on the performance of MULTICOM-CONSTRUCT. The A, C, and E show the plots of MAE_16 of long-range distance predictions vs the common logarithm of Neff of DeepAln, DeepMSA, and HHblits_BFD and their correlation coefficients, respectively. The B, D, and F show the plots of the long-range top L/2 contact precisions vs the common logarithm of Neff of MSA generated by DeepAln, DeepMSA, and HHblits_BFD and their correlation coefficients, respectively.

The correlation coefficients between top L/2 precision and the common logarithm of Neff for DeepAln and DeepMSA are 0.417 and 0.462, respectively. The correlation between the two is not very strong, mainly because some targets have a large Neff but low prediction accuracy due to the existence of the false-positive sequences in MSAs. 10 (or 9) out of 31 targets that have a Neff > 10 for DeepAln (or DeepMSA) have the precision of < 50%. Interestingly, the correlation coefficient between the top L/2 precision and the common logarithm Neff is 0.357 for the HHblits_BFD on all the 31 FM and FM/TBM CASP14 targets, which is even lower than DeepAln and DeepMSA. The correlation coefficients between the precision of the multi-class classification of distances and the logarithm of Neff are 0.373, 0.414, and 0.295 for DeepAln, DeepMSA, and HHblits_BFD, respectively, which is lower than the correlation for the binary contact prediction. The correlation coefficients between the MAE of multi-classification of distances and the common logarithm Neff are −0.488, −0.546, and −0.370 for the DeepAln, DeepMSA, and BFD, respectively. These results show that there is only a weak correlation between Neff and the accuracy of distance predictions for the three MSA generation pipelines (DeepAln, DeepMSA, Hblits_BFD), while the correlation is weakest for HHblits_BFD that generates the MSAs of the best quality.

Therefore, we conclude that both the quality and depth of MSAs impact the accuracy of distance predictions, and the depth measured by Neff is only a weak indicator of the accuracy of distance prediction. Indeed, there are some CASP14 targets (e.g., T1093) whose MSAs have a large Neff have low distance prediction accuracy. Different from the depth of MSAs that can be measured by a single quantity – Neff, the quality of MSA depends on alignment accuracy, and relationships between sequences (homologous or not) in MSA is hard to quantify.

### 3.3 Strong correlation between distance prediction accuracy and predicted probability scores and its application to select/combine predicted distance maps

According to the analysis above, different MSAs generated by different methods may work well on different sets of targets. Therefore, there is a need to find good metrics to select or combine MSAs or distance maps to improve prediction. However, Neff of MSAs has only a correlation with the accuracy of distance/contact prediction and therefore it cannot accurately select MSAs or predicted distance maps. In order to find better metrics to select MSAs and predicted distance amps, we calculate the correlation between the precision of top L/2 long-range contact predictions and the average probability of the top L/2 contact predictions (**Fig. 4**) The correlation between the two is 0.819. Moreover, the average probability also has a relatively strong correlation with the precision of multi-class classification of distances (correlation = 0.654) and the mean absolute error of the real-value distance prediction (correlation = −0.790). These correlations are much stronger than that between Neff and contact/distance prediction accuracy.

**Fig. 4.**
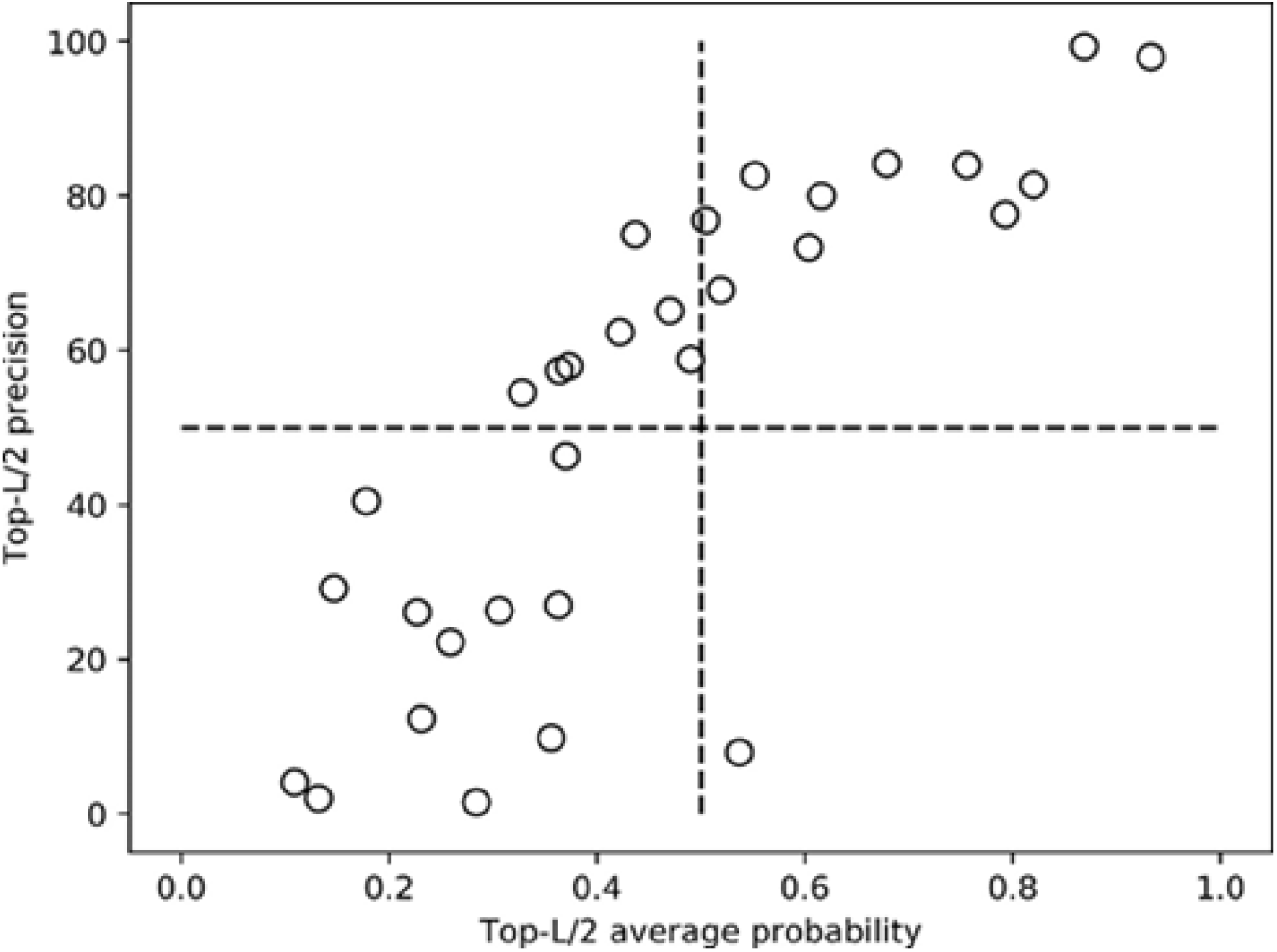
A plot of precisions of top L/2 long-range contact predictions against the average probabilities of the top L/2 predicted contacts. MULTICOM-CONSTRUCT with HHblits_BFD alignments were used to predict the distance maps.

The relatively strong correlation between the predicted contact probabilities and the accuracy of predicted distance maps provide a better approach to select distance maps predicted from different MSAs than Neff. To analyze the effectiveness of this approach for improving distance/contact predictions, we compare it with two approaches of combining MSAs or predicted distance maps: Combine_MSA_Map and Average_Map. Combine_MSA_Map merges the three MSAs generated by DeepAln, DeepMSA, and HHblits_BFD into one MSA file and uses CD-HIT (Li and Godzik, 2006) and HH-filter to do two rounds of redundancy filtering to generate a final MSA for MUlTICOM-CONSTRUCT to predict a distance map. Average_Map simply calculates the average of the distance maps predicted from the three MSAs as the final distance map prediction. We use Probability_Map to denote the approach of using the average probability of top L/2 long-range contact predictions to select a distance map from the three distance maps predicted from the three MSAs. Finally, Optimal_Map represents the ideal approach of always of selecting the most accurate distance map in terms of evaluation metric (top L/2 precision, top L precision, Precision_m, Recall_m, and MAE_16) from the three maps predicted from the three MSAs, which is the upper limit that any distance map combination or selection methods can reach.

**Table 4** reports the distance prediction results of using these approaches to select or combine the distance maps predicted from the three kinds of MSAs. Probability_Map works better than both Average_Map and Combine_MSA_Map in terms of almost all metrics and its performance is even close to Optimal_Map, indicating that the probability of top predicted contacts is a good metric to select distance maps predicted from different MSAs to improve distance prediction.

It is worth noting that Combine_MSA_Map performs worse than always selecting the distance maps predicted from the HHblits_BFD MSAs that works better than the MSAs of DeepAln and DeepMSA on average. The reason is that a simple combination of the MSAs from HHblits_BFD, DeepAln, and DeepMSA may introduce some noise (i.e. false positive – non-homologous sequences) into MSA, even though there are more sequences in the combined MSAs (higher depth).

**Table 4.** The performance of different methods of selecting/combining distance maps on 31 CASP14 full-length hard targets. The prediction distance maps were predicted by MULTICOM-CONSTRUCT with three different MSAs generation pipelines.

| Method | Top-L/2(%) | Top-L(%) | MAE_16(Å) | Precision_m(%) | Recall_m(%) |
| --- | --- | --- | --- | --- | --- |
| Combine_MSA_Map | 49.37 | 37.69 | 4.254±2.78 | 32.80 | 20.30 |
| Average_Map | 52.10 | 38.99 | 4.405±2.71 | 34.18 | 18.86 |
| Probability_Map | 52.25 | 39.27 | 4.078±2.77 | 33.38 | 21.44 |
| Optimal_Map | 52.5 | 40.7 | 4.010±2.64 | 36.65 | 21.58 |

### 3.4 Comparison of different feature sets on distance prediction

Each of the MULTICOM distance predictors uses four different sets of features derived from an MSA to predict distance maps and then average them as the final prediction from the MSA to improve the accuracy and stability of prediction. **Table 5** summarizes the distance prediction performance of four different feature sets using MULTICOM-CONSTRUCT with HHblits_BFD alignments on 37 FM and FM/TBM domains in comparison with the ensemble approach of averaging the four predicted distance maps from the four sets of features as the prediction. The ensemble approach performs better than using each feature set alone in terms of all evaluation metrics. Its mean precision of top L/2 long-range contacts is 50.18%, which is 3.33, 3.34, 3.47, and 6.19 percentage points higher than COV_set, PLM_set, PRE_set, and OTHER_set, respectively. The mean absolute error of the ensemble approach is 3.95 Å, lower than all the four feature sets. Also, the precision of the multi-class classification is 33.55%, higher than each feature set.

**Table 5.**
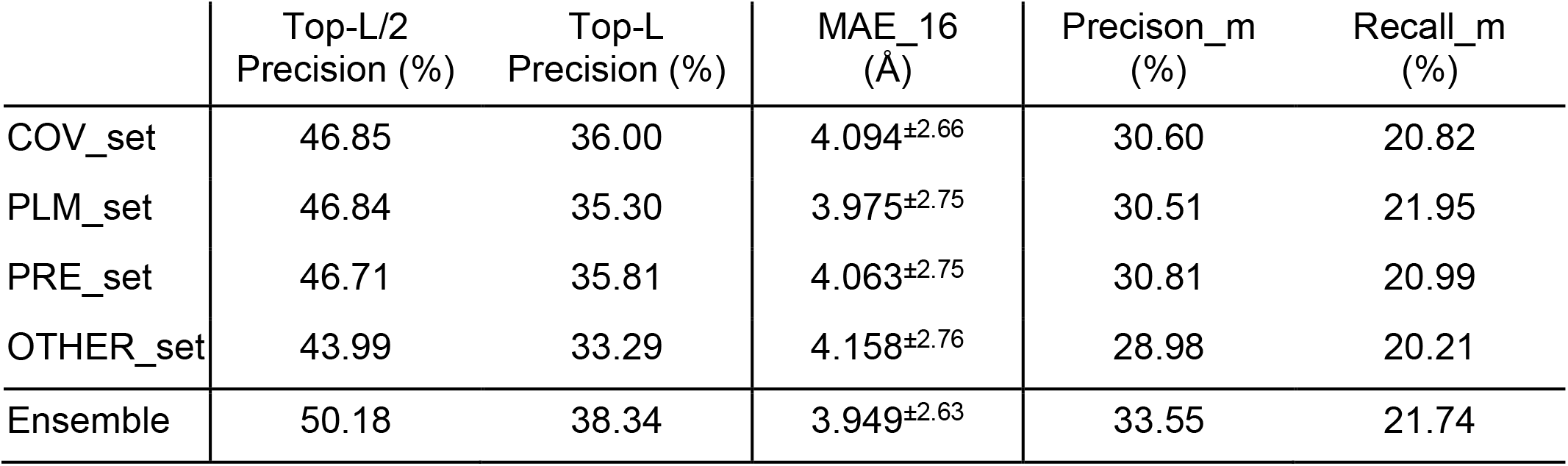
The performance of distance prediction on the 37 FM and FM/TBM domains with each of four feature sets: COV_set, PLM_set, PRE_set, and OTHER_set in comparison with the ensemble approach

Although the average performance of the ensemble approach is better, it does not perform best on every individual target. **Fig. 7** compares the max long-range top L/2 contact precision (diamond shape), average long-range top L/2 contact precision of four feature sets (square shape), and the long-range top L/2 contact precision of the ensemble approach (triangle shape). The results of the ensemble are not as good as the results of the best single feature set, especially for the target T1040-D1, T1047s1-D1, T1049-D1, T1082-D1, and T1096-D2 which are marked by red arrows. The gaps between the max precision of four feature sets and the precision of the ensemble approach on these targets are all greater than 8%, suggesting that there is still some room for improving the combination of features.

**Fig. 7.**
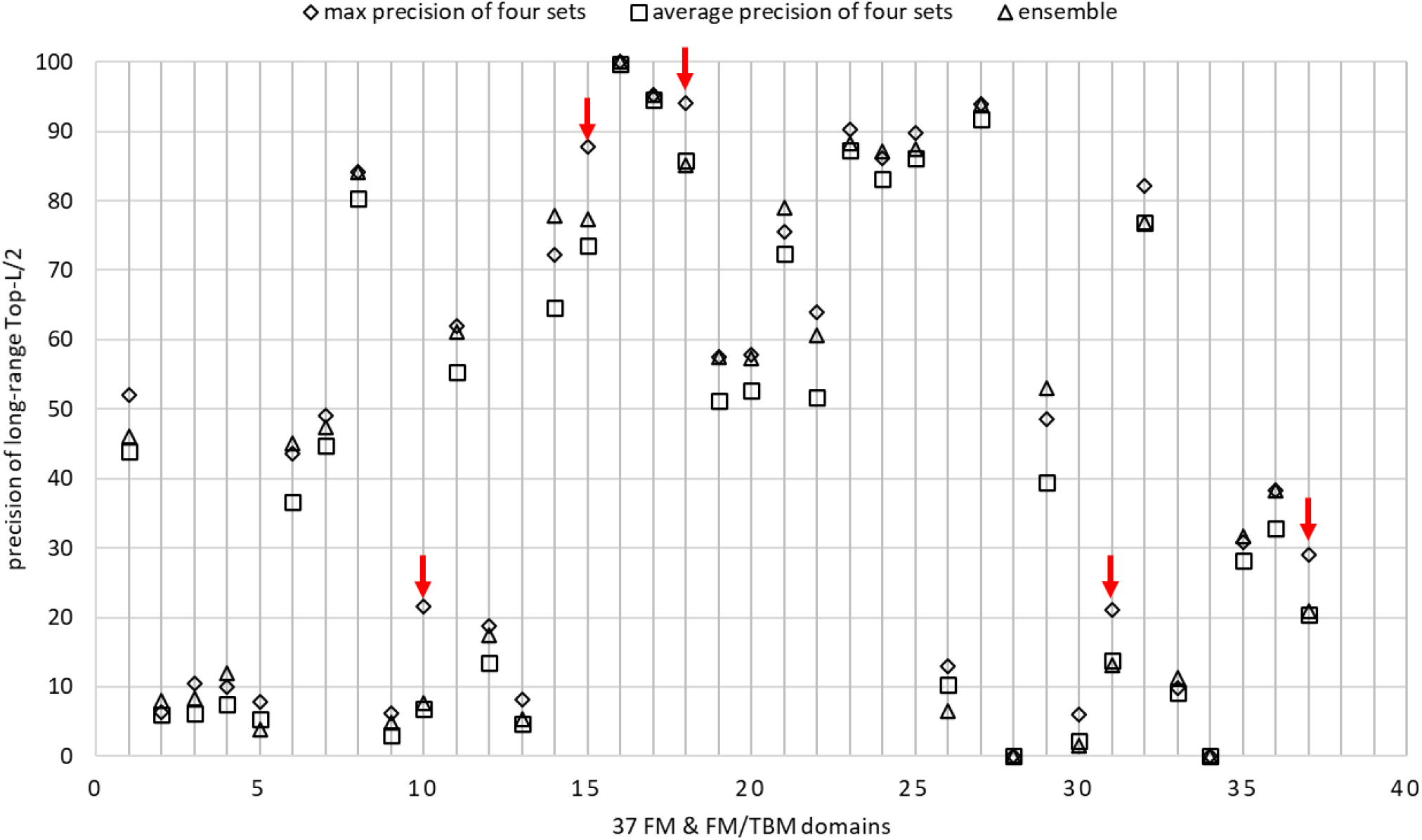
The comparison between the max top L/2 precision of four feature sets, the average precision of four feature sets, and the precision of the ensemble approach on 37 FM and FM/TBM CASP14 domains.

As a special case, **Fig. 8** illustrates the top L/2 long-range contacts of T1047s1-D1 predicted by the ensemble approach and from the PRE_Set in comparison with true contacts. The ensemble approach predicted more false positives marked in the eclipse than the PRE_Set.

**Fig. 8.**
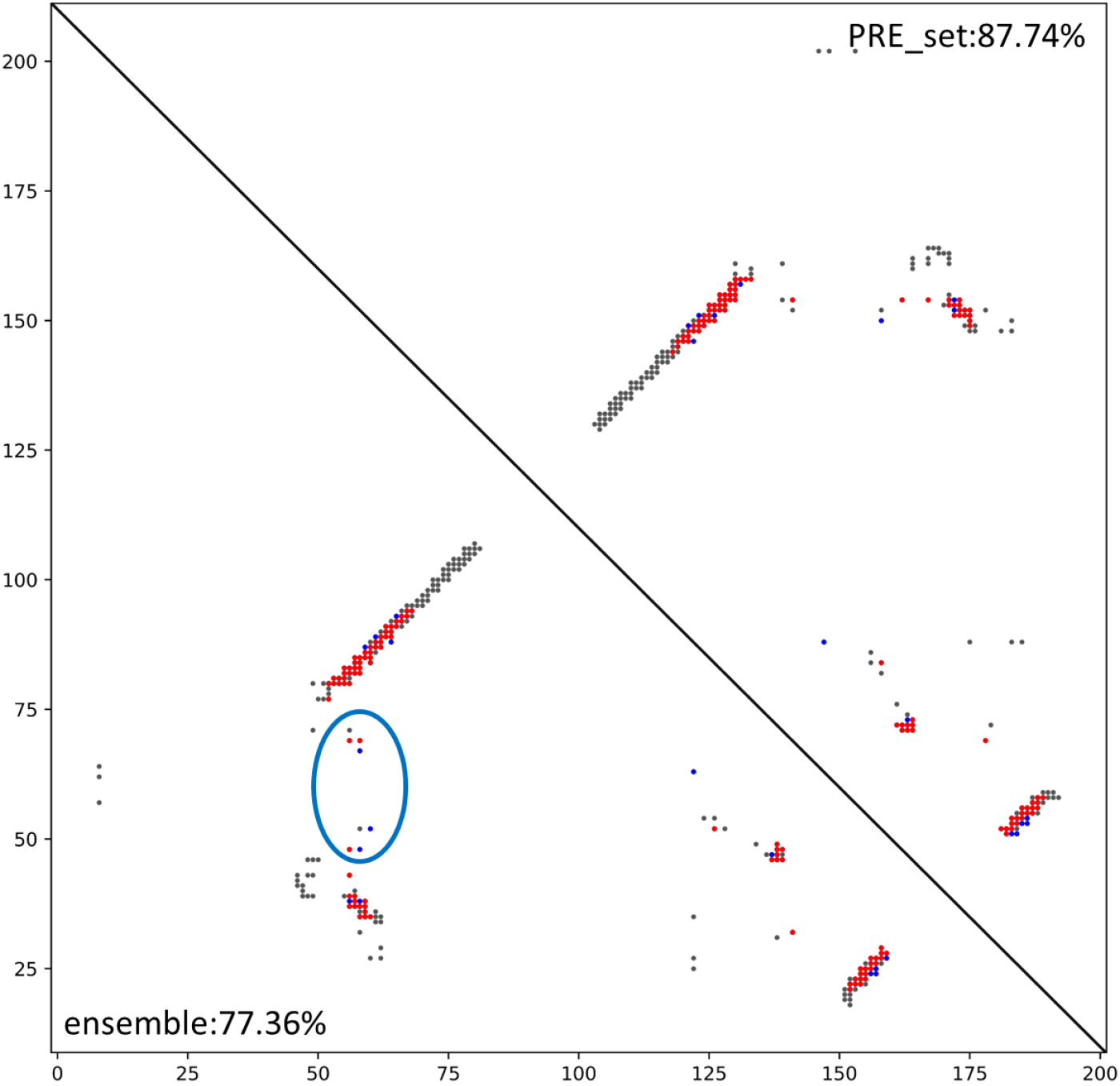
Top L/2 long-range contacts predicted from PRE_set (upper triangle: red dots: correct predictions and blue dots: incorrect predictions) and the ensemble (lower triangle) for T1047s1-D1 mapped onto the true contacts (gray dots). The blue eclipse circles some false positives predicted by the ensemble approach.

After CASP14, we tried to ensemble the distance prediction of multiple deep learning models trained on a single feature set and found that the integration of the results of multiple models can improve the stability and accuracy of the prediction. **Table 6** shows the comparison of a single deep learning model and the ensemble of four deep learning models that were trained on the COV_set and based on the approach similar to MULTICOM_CONSTRUCT. The performance of the ensemble of the four deep learning models using COV_set on CASP14 37 FM and FM/TBM domains is better than the single model in terms of all the evaluation metrics. The same phenomenon is also observed for the other three feature sets. Moreover, the ensemble of the four ensembles of the four feature sets obtains the long-range top L/2 contact prediction precision of 51.80%, the mean absolute error of 2.687Å, and the multi-classification precision of 34.17%, which is better than the ensemble of four single deep learning models trained on the four feature sets (i.e., 50.18%, 3.949Å, and 33.55% in **Table 5**).

**Table 6.** The performance of distance prediction of the single deep learning model and an ensemble of four deep learning models using the COV_set features on 37 CASP14 FM and FM/TBM domains.

| | Top-L/2<br>Precision<br>(%) | Top-L<br>Precision<br>(%) | MAE_16($\text{\AA}$) | Precision_m<br>(%) | Recall_m<br>(%) |
| --- | --- | --- | --- | --- | --- |
| COV_set<br>(single model) | 46.85 | 36.00 | $4.094 \pm 2.66$ | 30.60 | 20.82 |
| COV_set<br>(ensemble) | 49.20 | 37.84 | $2.638 \pm 2.11$ | 33.69 | 21.89 |

### 3.5 Impact of the size of the training dataset on prediction accuracy

We investigated the impact of the size of training datasets on the accuracy of protein distance prediction using the deep learning model of MULTICOM-CONSTRUCT on CASP14 37 FM and FM/TBM domains. MULTICOM-CONSTRUCT was trained on two datasets of different sizes. Dataset_1 introduced in DeepDist1 has 6463 proteins. Dataset_2 has 11034 proteins. The precision of top L/2 long-range contact predictions for the deep learning model trained on Dataset_2 is 50.18%, nearly 3% percentage point higher than on Dataset_1. A target-to-target comparison of mean absolute error (MAE) on 37 domains for the two models is shown in **Fig. 9**. On almost all the domains, the model trained on Dataset_2 has a lower MAE than that on Dataset_1. In some cases, such as T1038-D2, the difference is substantial.

**Fig. 9.**
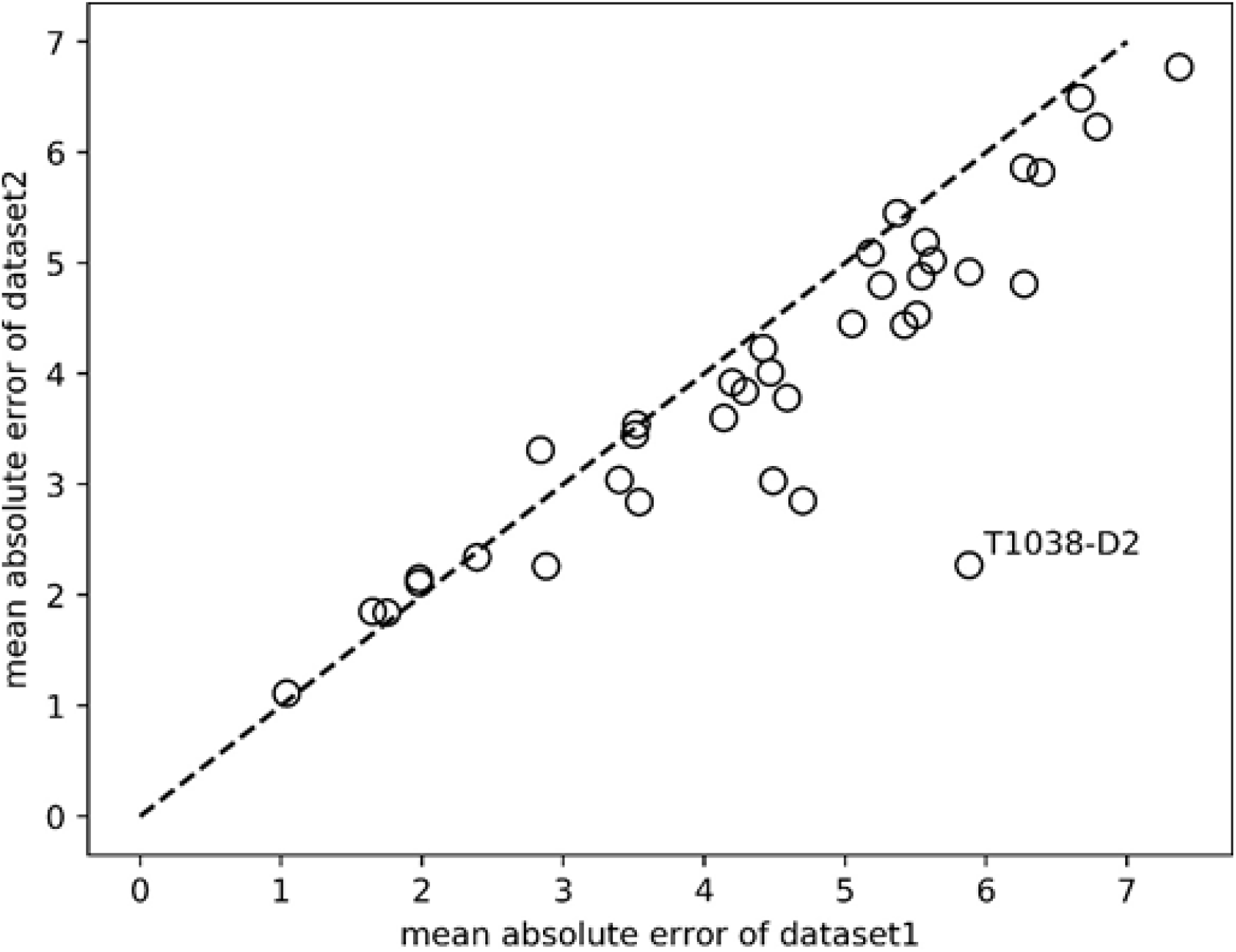
Comparison of the mean absolute error (MAE_16) of long-rang distance predictions of the two deep learning models trained Dataset_1 (small) and Dataset_2 (large).

The comparison between the distance maps predicted by the deep learning models trained on Dataset_1 and Dataset_2 and the true distance map of T1038-D2 is illustrated in **Fig. 10**. The distance map predicted by the model trained on Dataset_2 is very similar to the true distance map, but the distance map predicted by the model trained on Dataset_1 is very different.

**Fig. 10.**
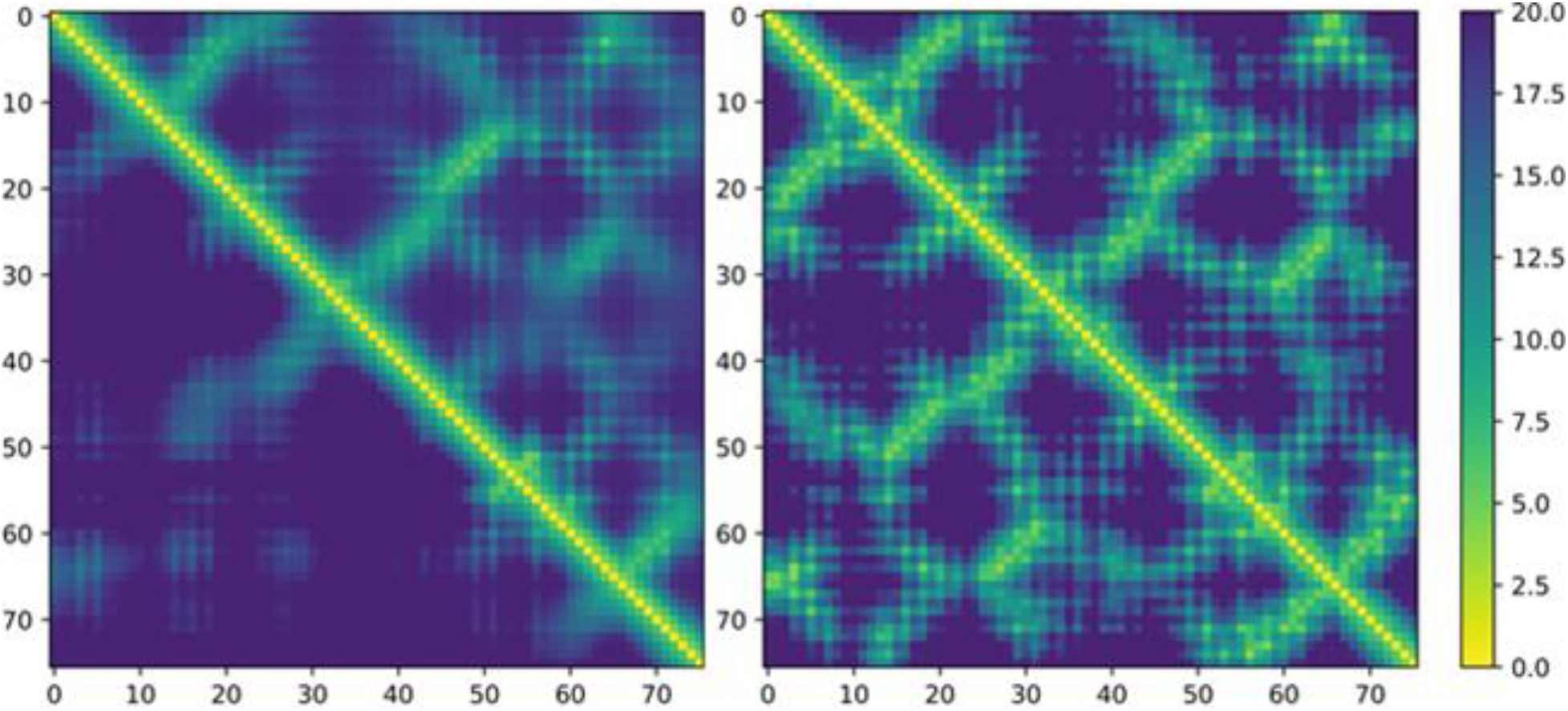
Comparison of predicted and true distance maps of T1038-D2. The upper triangle of the subfigure on the left is the distance map predicted by the model trained on Dataset_2 and the lower-triangle is the distance map predicted by the model trained on Dataset_1. The subfigure on the right is the true distance map. The distance map predicted by the model trained on Dataset_2 is much more similar to the true distance map than the model on Dataset_1.

### 3.6 The study of good and bad CASP14 cases

The MULTICOM distance predictors performed very well on T1052-D3. The average precision of top L/2 long-range contact predictions of MULTICOM predictors is close to 100%, while the average top L/2 precision of all CASP14 server predictors is 58.13%. T1052 is a multi-domain protein that has 832 amino acids, Neff of the MSA of the full-length T1052 is less than 15. The domain parsing program of MULTICOM predictors was able to identify a hard modeling region [590, 688] covering the range ([589, 668]) of the third domain of the target (T01052-D3) well. The sequence of the region was used to search against the sequence databases to build deeper MSAs to predict distance maps for the region. The distance maps predicted for the regions were combined with the full-length distance maps as in DeepDist (Wu, et al., 2020). This domain-based distance map prediction substantially increased the quality of the distance prediction for T1052-D3.

**Fig.11** compares the domain-based distance map prediction and the full-length distance map prediction made by MULTICOM-CONSTRUCT with the true distance map of T1052-D3. The domain-based distance map prediction is much better and clearer than the full-length distance map prediction for T1052-D3. The results show that good domain parsing can improve the quality of MSAs and therefore the quality of distance prediction.

**Fig. 11.**
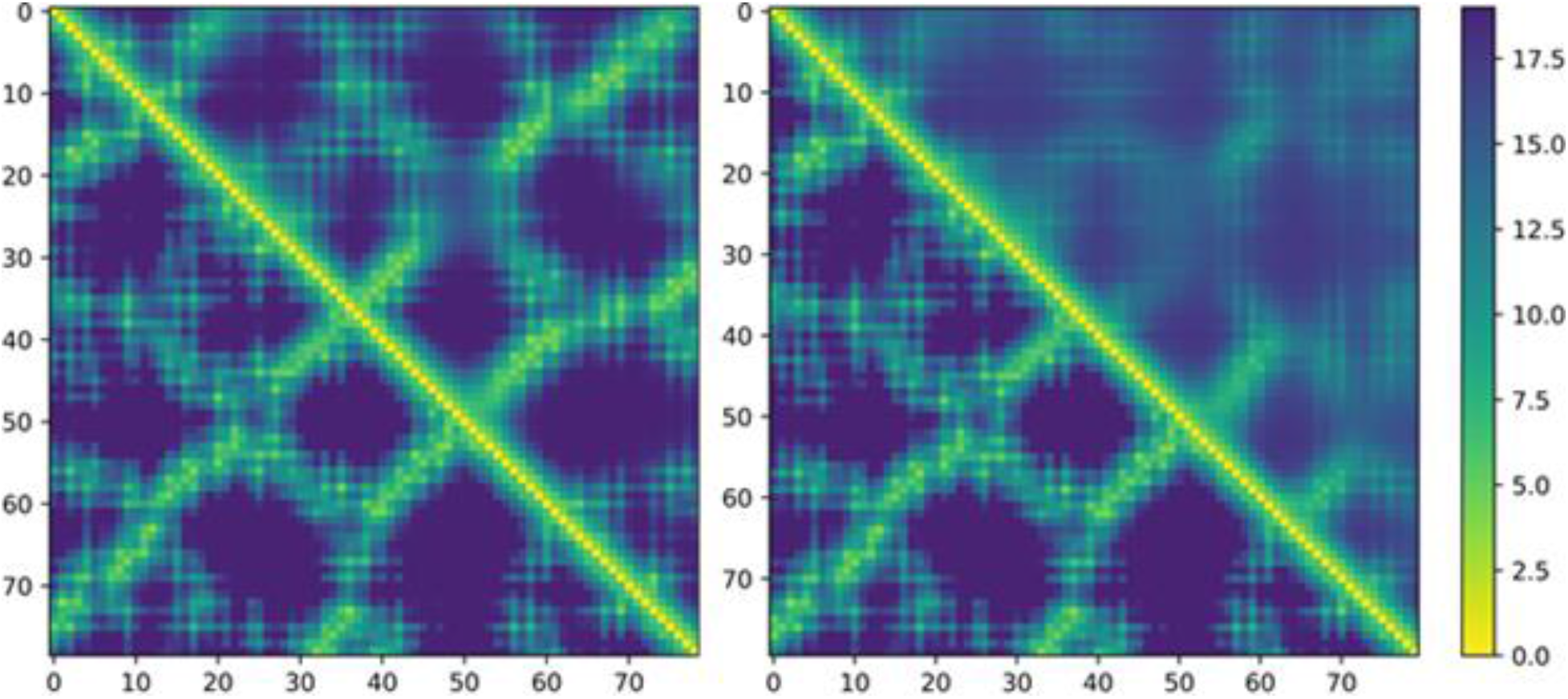
Comparison of the domain-based distance prediction and the full-length distance prediction with true distance map of T1052-D3. In the subfigure on the left, the upper triangle denotes the domain-based distance prediction, and the lower triangle the true distance map. In the figure on the right, the upper triangle denotes the full-length distance prediction, and the lower triangle the true distance map. The patterns in the domain-based distance prediction map are much clear and closer to the true distance map than the full-length distance prediction map.

Usually, the poor prediction of protein distances is due to a lack of effective sequences in MSAs (e.g., lower Neff on T1029, T1033, T1043, T1064) to generate good input features. The deep learning predictors cannot effectively extract distance patterns from them. However, in some cases, MSAs have high Neff, but the accuracy of the distance prediction is still very low. For instance, the Neff of the MSAs generated by DeepAln for T1093 is 689.36 and that generated by DeepMSA is 425.12, which are high values. However, all of the MULTICOM predictors got 0% top L/2 contact prediction precision, even the domain of the target can be reasonably identified. **Fig. 12** compares the distance maps predicted by four different approaches with the ground truth: (1) the distance map predicted from MSAs generated from DeepAln and DeepMSA with the predicted domain information (our original CASP14 submission, denoted as Original_dm), (2) the distance map predicted from MSA generated by HHblits_BFD without utilizing the domain information (denoted as BFD_full), (3) the distance map predicted from MSA generated by HHblits_BFD with the predicted domain information (denoted as BFD_dm). All these four distance maps above were predicted by MULTICOM-CONSTRUCT to ensure consistency.

**Fig. 12.**
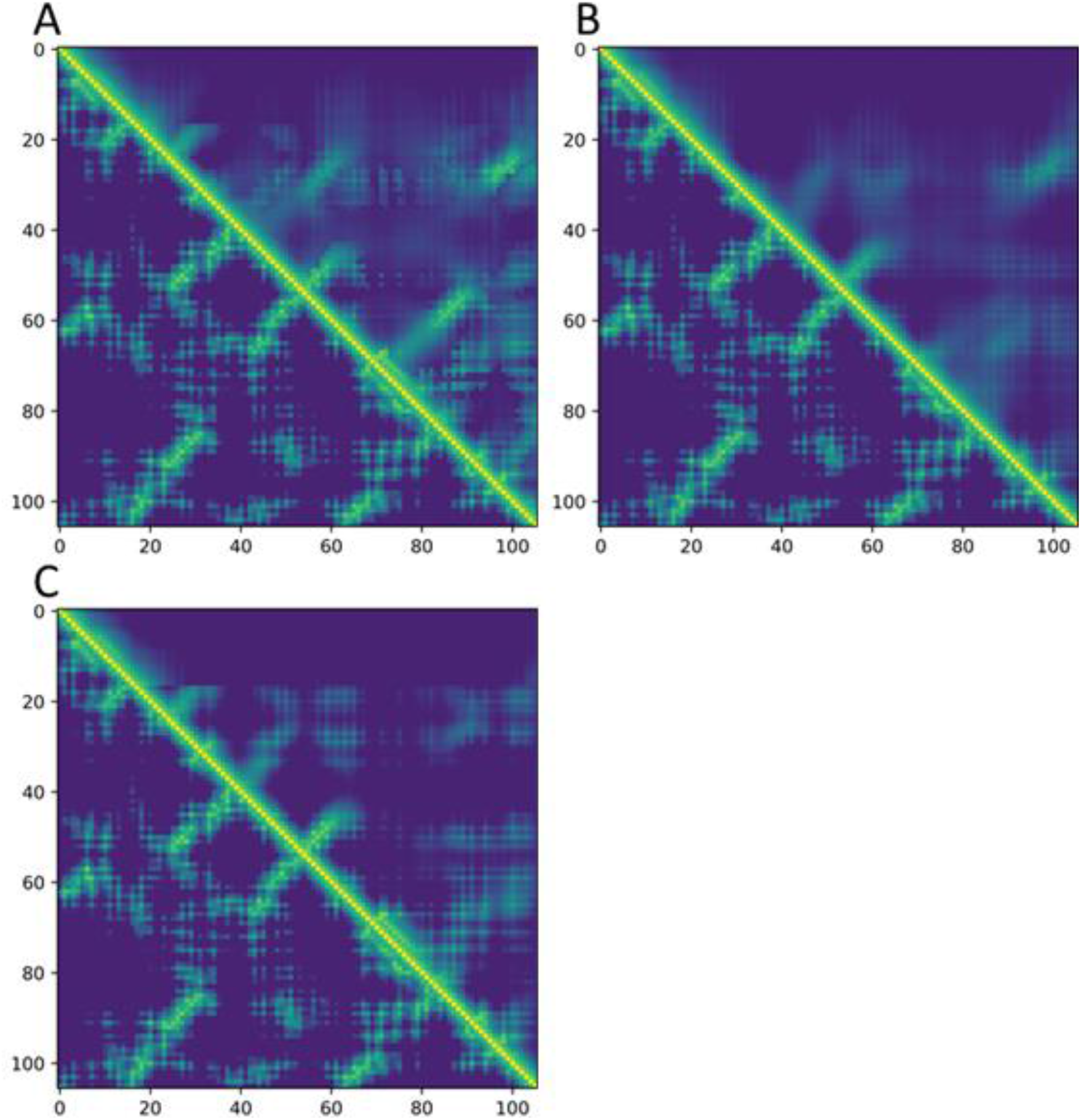
(A) The distanced map predicted from MSAs generated by DeepAln and DeepMSA with predicted domain information (upper triangle) versus the true distance map (lower triangle), (B) The distance map predicted from the HHblits_BFD MSA without domain information (upper triangle) versus true distance map (lower triangle), (C). The predicted distance map from the HHblits_BFD MSA with predicted domain information (upper triangle) versus the true distance map (lower triangle).

It can be seen that although MSAs generated by DeepAln and DeepMSA have a lot of sequences, most of them are false-positive positives leading to the prediction of many false-positive contact predictions (Fig. 12A). In **Fig. 12B**, the distance map predicted from the HHblits_BFD MSA without using predicted domain information is somewhat better, indicating that HHblits_BFD MSA (Neff = 133.0) has the better quality than MSAs of DeepAln and DeepMSA. If the predicted domain information is used, the distance prediction predicted from HHblits_BFD MSA is further improved in **Fig. 12C**, even though the Neff of the HHblits_BFD MSA for the domain is only 15, which is much lower than MSAs of DeepAln and DeepMSA. The long-range top L/2 contact prediction precision, the MAE of long-range distance prediction less than 16 Å, and the precision of multi-classification of distances using the different approaches for this domain are reported in **Table 7**. This case shows the quality of MSAs is important for distance prediction, and Neff is not always a good indicator of the quality of MSAs when there are false positives in MSAs.

**Table 7.**
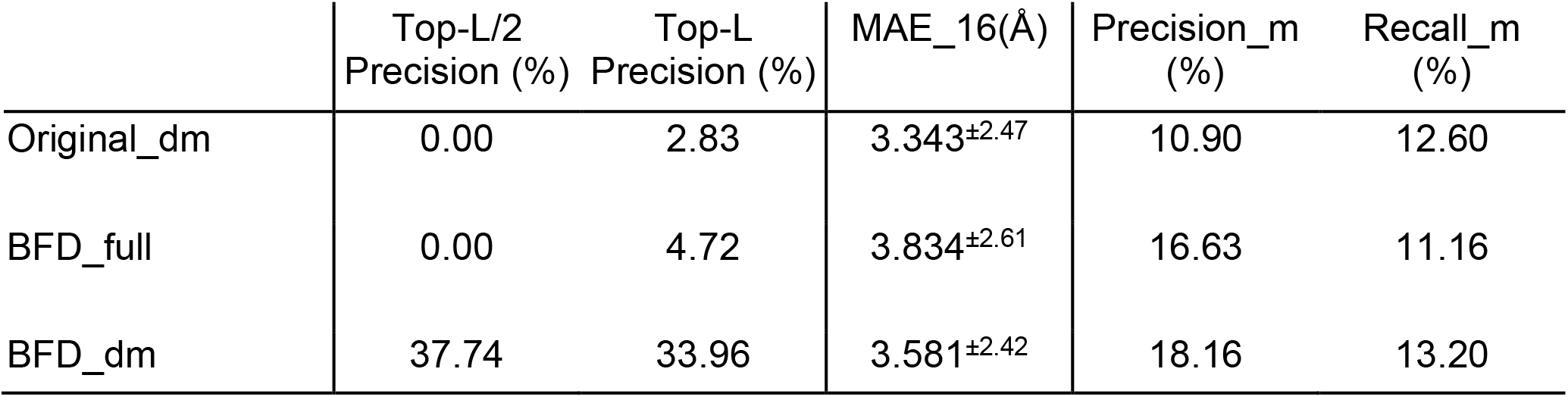
The accuracy of the distance prediction of using the three different approaches to generating MSAs for T1093.

|  | Top-L/2<br>Precision (%) | Top-L<br>Precision (%) | MAE_16(Å) | Precision_m<br>(%) | Recall_m<br>(%) |
| --- | --- | --- | --- | --- | --- |
| Original_dm | 0.00 | 2.83 | 3.343<sup>±2.47</sup> | 10.90 | 12.60 |
| BFD_full | 0.00 | 4.72 | 3.834<sup>±2.61</sup> | 16.63 | 11.16 |
| BFD_dm | 37.74 | 33.96 | 3.581<sup>±2.42</sup> | 18.16 | 13.20 |

## 4 Conclusion and future work

We developed several deep learning distance predators and rigorously benchmarked them in CASP14. The predictors performed reasonably well in the highly competitive CASP14 experiment. The results demonstrate that MSAs generated from different alignment methods on different databases for distance prediction have different quality. The MSAs generated by HHblits on the BFD database leads to the most accurate distance prediction, but different MSAs are still complementary and can be combined to improve distance prediction. However, the number of effective sequences of MSAs has only a weak correlation with the quality of MSA and therefore is not a strong factor of the quality of MSAs and the accuracy of the distance maps predicted from them because of the frequent existence of false positives in MSAs. In contrast, the predicted probabilities of top long-range contact predictions have a strong correlation with the accuracy of distance map predictions, and therefore is a better metric to select or combine predicted distance maps to improve distance prediction. Moreover, we show that the distance maps predicted from different features generated from the same MSA are also complementary and can be integrated to improve prediction accuracy. Finally, using larger training datasets to train deep learning models, ensembling multiple deep learning models, or applying domain predictions to MSA generation of some multi-domain targets can also improve the accuracy of the distance prediction.

## Acknowledgments

The project is partially supported by two NSF grants (DBI 1759934 and IIS1763246), one NIH grant (GM093123), two DOE grants (DE-SC0020400 and DE-SC0021303), and the computing allocation on the Summit supercomputer provided by Oak Ridge Leadership Computing Facility (Project ID: BIF132).

